# Biopurification system as a source of pesticide-tolerant bacteria able to degrade the commonly used pesticides chlorpyrifos and iprodione

**DOI:** 10.1101/489252

**Authors:** M. Cristina Diez, Claudio Lamilla, Bárbara Leiva, Marcela Levio, Pamela Donoso-Piñol, Gabriela Briceño, Heidi Schalchli, Felipe Gallardo

## Abstract

Intensive use of pesticides applied simultaneously in field to improve the effectiveness of pest control increase the environmental contamination, affecting the soil and water quality. Some of the commonly used pesticides are the insecticide chlorpyrifos and the fungicide iprodione; being thus critically essential to develop bioremediation methods to remove these contaminants by tolerant-bacteria. In this study we selected and characterized different pesticides-tolerant bacteria isolated from a biomixture of a biopurification system that had received continuous applications of a mixture of the pesticides chlorpyrifos and iprodione. Out of the 10 isolated bacterial colonies, only six strains presented adequate growth in presence of the both pesticides at 100 mg L^−1^. Biochemical and enzymatic characterization using API ZYM showed that all isolates (100%) were positive for esterase, leucine aminopeptidase, acid phosphatase, and naphthol-AS-BI-phosphohydrolase. According to the molecular level study of the 16S ribosomal gene and MALDI TOF/TOF MS, it was possible to determine that the isolated bacteria belong to the genera *Pseudomonas*, *Rhodococcus* and *Achromobacter*. Bacterial growth decreased proportionally (R^2^ > 0.96) as been as both pesticide concentrations increased from 10 to 100 mg L^−1^. *Achromobacter* sp. strain C1 showed the best chlorpyrifos removal (between 56–29%) after 120 h of incubation. On the other hand, the highest iprodione removal (between 91.2–98.9%) was observed for the *Pseudomonas* sp. strain C9, which was not detected after 48 h of incubation. According with their identification and ability to remove the contaminants, *Achromobacter* sp. strain C1 and *Pseudomonas* sp. strain C9 appear as promising microorganisms for their use in the treatment of matrices contaminated with chlorpyrifos, iprodione or their mixture. The results of this study will help to improve current technologies for the biodegradation of this commonly used insecticide and fungicide, in order to give a response to the problem of contamination by pesticides.

## Introduction

Different pesticides are applied simultaneously in the field to improve the effectivity of pest control, thus increasing environmental contamination and affecting the soil and water quality [1,2]. Toward minimizing pesticide point-source contamination, a preventative technology of biopurification named biobed was introduced and implemented in Sweden in the 90s by Torstensson and Castillo [3] to reduce the risk of water resources contamination. Pesticides removal by this biopurification technology is based on the adsorption and degradation capacity of an organic biologically active matrix (biomixture) composed of top soil, peat, and lignocellulosic material and a vegetal layer [4,5]. The biopurification system is highly efficient in pesticide removal, achieving high degradation of different pesticides commonly applied in farms, even after repeated applications [6,7]. The microbial communities are considered a key factor to control the depuration capacity of the biopurification system, and knowledge of the biological activity occurring in the biomixture is relevant for understanding pesticide degradation and to optimize their degradation [7–9]. In this respect, some studies have correlated pesticides degradation in the biomixture with microbial activities such as phenoloxidase activity [10–12], respiration rate [13], and microbial community changes [6,13–15].

A genotypically and phenotypically versatile microbial community can degrade different pesticide residues at different concentrations in a biomixture [6]. A greater bacterial diversity compared to fungal diversity has been reported, such that bacterial diversity increased throughout the biopurification system as affected by pesticide exposure [16].

It is well known that microorganisms are responsible for the degradation of pesticides in soils [17,18]. This is due to the extensive use of these compounds in agricultural soils, which has induced mechanisms of genetic adaptation in microorganisms. These genetic adaptations have led to the synthesis of enzymes that oxidize, hydrolyze, and hydroxylate pesticides, allowing them to use pesticides as the sole source of carbon, nitrogen, sulfur, or phosphorus and facilitating the elimination of the compound’s toxicity [19]. Further, active microbial populations develop in the soil with the ability to degrade persistent compounds after repeated pesticide application in the same field for a certain number of years [20].

Although bacterial species play important roles in the transformation of pesticides, the complete mineralization of pesticide residues is more likely to occur with mixed populations than individual microorganisms [21]. Fungi have also been reported as good pesticide degraders. A new fungal strain Hu-01 isolated from an activated sludge sample from an aerobic chlorpyrifos-manufacturing wastewater treatment plant, identified as *Cladosporium cladosporioides*, showed high chlorpyrifos degradation activity [22].

Among bacteria responsible for the degradation of pesticides, the genera *Streptomyces*, *Arthrobacter*, and *Achromobacter* have been isolated from contaminated soil and soil with historical application and studied due to their great capacity to degrade various pesticide residues, including CHL and IPR [23–26].

Campos *et al*. [25] reported two strains, *Arthrobacter* sp. strain C1 and *Achromobacter* sp. strain C2 isolated from soil, which are able to transform IPR and its degradation metabolite (3,5-dichloraniline) in different culture media. The degradation of IPR by the *Arthrobacter* strain C1 proceeded rapidly in all media with complete degradation observed within 8 and 24 h of culture, and this strain maintained its degrading capacity in a wide range of temperatures and pH. In contrast, *Achromobacter* sp. strain C2 was only able to slowly co-metabolize IPR. Additionally, metabolic intermediates 3,5-dichlorophenyl-carboxamide and 3,5-dichlorophenylurea-acetate in the metabolic IPR pathway, produced by these soil bacteria and their combination, were reported by Campos *et al*. [27].

Briceño *et al*. [23] reported two *Actinobacteria* isolated from an agricultural soil that had received continuous applications of CHL, which were able to rapidly degrade CHL with approximately 90% degradation after 24 h of incubation. These two strains were identified as *Streptomyces* sp. (AC5 and AC7 strains). Despite the high CHL degradation by both strains, a different behavior was observed when its main metabolite, 3,5,6-trichloro-2-pyridinol (TCP), was analyzed. A lower concentration of TCP (0.46 mg L^−1^) was produced by *Streptomyces* sp. strain AC5, and its concentration decreased as a function of time, as the TCP produced was 10 times lower compared to that produced by *Streptomyces* sp. AC7 strain (4.32 mg L^−1^).

Further, several studies have reported the potential of indigenous microbial consortia isolated from contaminated soils to degrade different pesticides and pesticide mixtures. In this context, Fuentes *et al*. [28] reported a *Streptomyces* sp. consortium able to remove an organochlorine pesticide mixture composed of lindane, methoxychlor, and chlordane. Recently, mixed cultures of the fungus *Trametes versicolor* and *Streptomyces* spp. were used to inoculate different biomixtures based on their previously demonstrated ligninolytic and pesticide-degrading activities [21]. The authors demonstrated that the consortium improved lindane dissipation (81–87%) or removal at 66 d of incubation in different biomixtures, decreasing the lindane half-life to an average of 24 d, 6-fold less than the T50 of lindane in soils. In addition, Briceño *et al*. [29] reported for the first time the removal of the organophosphorus pesticides mixture composed of CHL and diazinon from different environmental matrices (liquid medium, soil, and a biobed biomixture) by a *Streptomyces* mixed culture.

The previous studies mentioned have reported the ability of selected bacteria isolated from pesticide-contaminated soils to remove pesticides. However, the isolation and characterization of pesticide-degrading microorganisms from a biopurification system used for pesticide treatment have been scarcely studied. Therefore, the goal of this study was to select and characterize bacterial species isolated from a biopurification system and with the ability to degrade the fungicide IPR and the insecticide CHL.

## Materials and Methods

### Pesticides and culture media

Analytical grade (99%) iprodione (IPR), 3,5-dichloroaniline (3,5-DCA), chlorpyrifos (CHL), and 3,5,6-trichoro-2-pyridinol (TCP) for chromatographic analyses by HPLC were purchased from Sigma-Aldrich (St. Louis, MO). The stock solutions (1000 mg L^−1^) in acetone were sterilized by filtration through 0.22-μm pore-size membranes. For degradation assays, formulated commercial CHL (Troya 4EC) and IPR (Rovral 50 WP) were purchased from Agan Chemicals Manufacturers Ltd. The characteristics of the commercial products are shown in Table 1. Commercial products were prepared individually in a stock solution of 10,000 mg L^−1^ in methanol, filtered through a 0.22-mm PTFE filter, and then stored at 4 °C until their use. All other chemicals and solvents were of analytical reagent grade (Merck).

**Table 1.** Physicochemical characterization for the tested commercial pesticides.

| Pesticide | Commercial product | Concentration | Kind | Chemical formula | Water solubility (mg L<sup>-1</sup>) | Molecular weight (g mol<sup>-1</sup>) | T<sub>1/2</sub> (d) | GUS | K<sub>oc</sub> |
| --- | --- | --- | --- | --- | --- | --- | --- | --- | --- |
| Chlorpyrifos | Troya 4 EC | 480 g L<sup>-1</sup> | Insecticide | C<sub>9</sub>H<sub>11</sub>Cl<sub>3</sub>NO<sub>3</sub>PS | 1.05 | 350.58 | 50 | 0.17 | 8151 |
| Iprodione | Rovral 50 WP | 500 g kg<sup>-1</sup> | Fungicide | C<sub>13</sub>H<sub>13</sub>Cl<sub>2</sub>N<sub>3</sub>O<sub>3</sub> | 6.8 | 330.17 | 36.2 | 0.58 | 700 |

Solubility in water at 20 °C; T<sub>1/2</sub>: Time half-life, GUS: Groundwater Ubiquity Score; K<sub>oc</sub>: Adsorption coefficient.

Mineral salts medium (MSM) broth containing (per L) 1.6 g K_2_HPO_4_, 0.4 g KH_2_PO_4_, 0.2 g MgSO_4_ · 7H_2_O, 0.1 g NaCl, 0.02 g CaCl_2_, and 1 mL salt stock solution (2.0 g boric acid, 1.8 g MnSO_4_ · H_2_O, 0.2 g ZnSO_4_, 0.1 g CuSO_4_, 0.25 g Na_2_MoO_4_, 1000 mL distilled water) was used for pesticide degradation assay. The initial pH of the medium was adjusted to 7.0 prior to sterilization by autoclaving (121 °C for 20 min). Subsequently, cycloheximide (0.05 g L^−1^) was added to avoid fungal contamination. Luria Bertani (LB) broth containing (per L) 5.0 g NaCl, 2.5 g yeast extract, and 10.0 g casein peptone was used for routine cultivation of the isolated bacteria. The pH of LB was adjusted to 7.0 prior to autoclaving. Plate count agar (PCA) containing (per L) 5.0 g tryptone, 2.5 g yeast extract, 1.0 g glucose, and 15.0 g agar-agar was adjusted to pH 7.2 prior to sterilization, and 0.05 g cycloheximide was added to avoid fungal contamination. Finally, R2A agar containing (per L) 0.5 g casein acid hydrolysate, 0.5 g yeast extract, 0.5 proteose peptone, 0.5 g dextrose, 0.5 g soluble starch, 0.3 g K_2_HPO_4_, 0.024 g MgSO_4_, 0.3 g sodium pyruvate, and 15 g agar, pH 7.2 was used for strains biochemical characterization.

### Biopurification system for isolation of pesticide-tolerant bacteria

Pesticide-tolerant bacteria were isolated from a biopurification system (BPS) used during the last three years for pesticide treatment (CHL and IPR at 50 mg kg^−1^ a.i. each) with re-application every 30 days 1/1/0001 12:00:00 AM. The BPS consisted of a plastic tank of 1 m^3^ capacity packed with 125 kg of biomixture (dry weight) (bulk density (ρ) 0.29 g mL^−1^), which reached a height of 60 cm. The biomixture used in BPS was prepared with top soil, commercial peat, and wheat straw in a proportion of 1:1:2 (v v^−1^), and humidity was maintained at about 65–70% of water holding capacity (WHC) by addition of tap water.

For strain isolation, biomixture subsamples were collected from different parts of the BPS, and a composite sample (500 g) was stored at 4 °C for no longer than 12 hours. Microorganisms in the biomixture were counted using the serial dilution method. For this, 10 g biomixture was added to 90 mL saline solution (0.9%), and the suspension was shaken vigorously. Subsequently, 150-µL aliquots of each dilution were inoculated on Petri dishes containing PCA medium. Incubation was performed at 28 ± 2 °C for 48 h, following which, the colonies formed were counted.

Pesticide-tolerant bacteria were isolated by placing 10 g of the biomixture in sterile 250-mL Erlenmeyer flasks containing 90 mL of MSM broth supplemented with CHL plus IPR at 10 mg L^−1^ a.i. each). Flasks were incubated for 7 days at 28 ± 2 °C and 130 rpm with constant shaking in the dark. After this period, decimal dilutions from 1 × 10^−1^ to 1 × 10^−4^ were prepared in order to obtain perfectly separated strains. For this, 65-µL aliquots of each dilution were inoculated on Petri dishes with 30 mL of PCA medium. Plates were incubated at 28 ± 2 °C for seven days, and morphology and coloration of the colonies were analyzed for bacterial selection. The bacterial strains were maintained on LB-glycerol (70/30%) medium slants at 4 °C, and they were filed at the Laboratory of Environmental Biotechnology in La Frontera University.

To examine the ability of the strains to grow in the presence of pesticides (CHL and IPR), a quantitative assay was performed. The study consisted of evaluating biomass growth in flasks containing 50 mL of LB broth supplemented with each pesticide at 10 mg L^−1^ concentration. The flasks were incubated at 28 ± 2 °C and 130 rpm under constant shaking during 48 h, and bacterial growth was measured by measuring the absorbance at 600 nm. Thereafter, absorbance values were converted to biomass dry weight (g L^−1^) using a calibration curve (R^2^ > 0.999).

### Characterization of selected pesticide-tolerant strains

The selected bacterial strains were characterized by a combination of phenotypic tests as described by Krishnapriya *et al*. [30], which are based mainly on colony morphology, Gram staining reaction, and colony pigmentation.

Visualization of pesticide-tolerant bacterial cells was performed using Scanning Electron Microscopy (SEM) with variable pressure (VP-SEM) and the instrument equipped with a STEM detector (Transmission Module) (SU-3500 Hitachi-Japan). The strain samples were obtained from a fresh 24-h culture in PCA. The bacterial colonies were washed three times with distilled water, and the pellet was re-suspended in LB medium at 0.5 McFarland with distilled sterile water and acetylchlorine (0.1%). A sample of 65 µL of each strain was placed in the equipment sampler and dried at 30°C, followed by microscopic observations.

The strains were subjected to biochemical characterization using the APIZYM kit (Biomerieux, France) according to the manufacturer’s instructions. This microbial identification system consists of 19 substrates in a microplate, which was incubated at 28 °C for up to 4 days. The enzyme activity was detected based on the intensity of color developed following the addition of reagents.

Moreover, extracellular hydrolyzing enzyme production was screened as described by Margesin *et al*. [31]. The presence of amylase, cellulase, lipase, protease, and gelatinase activity was tested on R2A agar supplemented with starch (0.4% w v^−1^), carboxymethylcellulose and trypan blue (0.4% and 0.01% w v^−1^), Tween 80 (1% v v^−1^), skim milk powder (0.4% w v^−1^), or gelatin (1% w v^−1^), respectively. The agar plates were prepared in triplicate. After 3–10 days at 15 °C, a positive reaction was observed when transparent zones around the colonies were directly visible or detected after precipitation or coloration of the non-degraded substrate. To reveal amylase and protease activities, the plates were stained with Lugol’s solution and Coomassie brilliant blue solution, respectively [31]. The assays were performed in triplicate plates.

### Bacterial identification by sequence analyses and MALDI-TOF/TOF MS

For identification of pesticide-tolerant bacteria, genomic DNA was extracted using the UltraClean^®^ Microbial DNA Isolation Kit (MOBIO, CA, USA) according to the manufacturer’s instructions. 16S rDNA was selectively amplified from genomic DNA by polymerase chain reaction (PCR) using universal primers 27F (5’-AGAGTTTGATCCTGGCTCAG-3’) and 1492R (5’-GGTTACCTTGTTACGACTT-3’), enabling the amplification of approximately 1.500 bp of the 16S rRNA gene. PCR amplification was performed in a Multigene Optimal Thermal Cycler (Labnet, USA) in 50 µL of PCR mix comprising 25 µL mix reaction buffer 2x (SapphireAmp Fast PCR Master Mix, Takara), 22 µL ultra-pure water, 1 µL of each primer (10 µM), and 1 µL of DNA. The temperature and cycling conditions were as follows: preheating at 94 ºC for 2 min; 30 cycles at 94 ºC for 1 min; 55 ºC for 1 min; 72 ºC for 1.5 min; and incubation at 72 ºC for 10 min. The presence of PCR products was assessed by electrophoresis on a 1% agarose gel stained with gel red. Sequencing was conducted using a dye Terminator Cycle Sequencing Kit and an ABI 3730XL DNA Sequencer (Applied Biosystems) by Macrogen (Korea). The nearest taxonomic group was identified by 16S rDNA nucleotide sequence BLASTN (http://www.ncbi.nlm.nih.gov/blast) using DDBJ/EMBL/GenBank nucleotide sequence databases. The phylogenetic affiliation of bacteria in GenBank was performed using MEGA7. For the MALDI-TOF/TOF MS analysis, samples of selected bacterial colonies were applied directly to the equipment sampler plate and coated with a saturated solution of α-cyano 4-hydroxy cinnamic acid diluted in 50% acetonitrile with 2.5% trifluoracetic acid. Mass spectra were obtained using a MALDI-TOF/TOF MS Autoflex Speed (Bruker Daltonics, Bremen, Germany) equipped with a smart beam laser source (334 nm). Analyses were performed in linear mode with positive polarity, acceleration voltage of 20 kV, and extraction with delay of 220 ns. Each spectrum was collected as an average of 1200 laser shots with enough energy to produce good spectra without saturation in the range of 2000 to 20,000 m/z. Analyses equipment was calibrated externally using the protein calibration standard I (Bruker Daltonics, Bremen, Germany) (insulin, ubiquitin, cytochrome C and myoglobin) with Flex Control 1.4 software (Bruker Daltonics, Bremen, Germany). The sample analyses were performed with the MALDI Biotyper Compass 4.1 software (Bruker Daltonics, Bremen, Germany) in the range of 3000–15000 m/z, compared with a library of 6509 spectra of bacterial identifications. According to the guidelines of the manufacturer, a score of ≥ 2 depicts identification to the species level, and an intermediate log score between < 2 and ≥ 1.7 for identification to the genus level. A dendrogram generated by MALDI Biotyper mass spectra was performed for all strains isolated after enrichment with CHL and IPR in liquid cultures.

### Pesticide degradation in liquid culture

A pesticide degradation assay was conducted with the six selected pesticide-tolerant and well-characterized strains. For obtaining inocula, the strains were re-activated on plate dishes with PCA medium and incubated for 24–48 h at 28 ± 2 °C. After that, the bacteria were initially grown at 28 ± 2 ºC for 48 h in Erlenmeyer flasks with LB broth supplemented with a mixture of 10 mg L^−1^ of IPR and CHL to acquire enough biomass for downstream inoculation. Biomass was collected by centrifugation (6000 rpm for 10 min), washed, and re-suspended using sterile NaCl (0.9%). The degradation experiments in liquid media were conducted in 100-mL flasks that contained 50 mL of LB broth supplemented with each individual pesticide at a concentration of 0, 10, 20, 50, and 100 mg L^−1^. Subsequently, the biomass inoculum was added at 1% (v v^−1^), and non-inoculated flasks were run as controls. The flasks were then incubated at 28 ± 2 °C on a rotary shaker at 130 rpm in the dark for 48 h and 120 h for IPR and CHL, respectively. Samples were taken at different times for analysis of biomass growth, residual pesticide concentrations (IPR and CHL), and metabolite (3,5-DCA and TCP) formation. For biomass growth and pesticide degradation, kinetics parameters were calculated.

### Analyses of pesticides and metabolites

One milliliter of each sample was taken directly from each flask, and centrifuged at 6500 rpm for 10 min. After that, 0.5 mL of the supernatant was diluted in 1 mL of acetonitrile grade HPLC. The sample was homogenized in a vortex and filtered by filter PTFE 0.22 μm before analysis of pesticide concentrations. Analysis was performed using a Merck Hitachi L-2130 pump equipped with a Rheodyne 7725 injector and a Merck Hitachi L-2455 diode array detector. Separation was achieved using a C18 column (Chromolit RP-8e, 4.6 μm × 100 mm). The mobile phase was 70% 1 mM ammonium acetate and 30% acetonitrile injected at a flow rate of 1 mL min^−1^. The column temperature was maintained at 30 ± 1 °C; the detector was set for data acquisition at 290 nm. Instrument calibrations and quantifications were performed against pure reference standards (0.01–10 mg L^−1^) for each pesticide. Average recoveries for the pesticide were: IPR, 92 ± 2.2%; CHL, 101 ± 0.7%. Limit of quantification (LOQ) was determined using the smallest concentration of the analyte in the test sample, which induced a signal that was ten times higher than the background noise level (CHL = 0.214 mg L^−1^ and IPR = 0.238 mg L^−1^). Limit of detection (LOD) was 0.081 for CHL and 0.089 for IPR.

### Kinetics and statistical analysis

Data obtained in the degradation assays were used to determine the specific growth rate for the exponential phase using the following equation: µmax = dx/dt × 1/x; where µ= specific growth rate (h^−1^), x = biomass concentration (g L^−1^), and t = time (h). The removal of IPR and CHL was described using the first-order kinetic model: ln Ct/C0 e-kt, where C0 is the amount of contaminant in the liquid medium at time zero, Ct is the amount of contaminant at time t, and k and t are the rate constant and degradation time in hours, respectively. The time at which the IPR and CHL concentrations in the liquid medium were reduced by 50% (T_1/2_) was calculated using the equation T_1/2_= ln (2)/k. In degradation study, simple correlation analysis was done to determine correlation between biomass growth and initial pesticide concentration.

Data were averaged and the standard deviation (SD) of the means was calculated. Removal percentage data were transformed using an angular transformation (arc sen √x/100) prior to statistical analysis. Post hoc analysis of differences in means of the assay data was conducted with the Tukey test (α=0.05). Statistical analyses were performed using SPSS statistical software version 17.

## Results

### Isolated bacteria from the biopurification system

Pesticide-tolerant bacteria were isolated from a biomixture used in a biopurification system, which in the last three years had been used to degrade a mixture of pesticides added repeatedly at a concentration of 50 mg L^−1^. To approximate the number of viable bacteria in the biomixture, a plate count test was performed. The results revealed 23 × 10^6^ UFC g^−1^ of biomixture in the PCA medium.

In the present study, 10 different types of bacterial colonies (strains C1–C10) isolated using PCA medium were obtained after enrichment with CHL plus IPR (10 mg L^−1^ a.i. each) from the biomixture. Out of the 10 bacterial colonies, only six strains presented adequate growth expressed as biomass concentration ≥ 1.0 g L^−1^ in LB broth, which had been supplemented with a mixture of CHL and IPR (10 mg L^−1^ a.i. each). Specifically, strains C4, C9, and C10 showed a biomass growth of > 2.0 g L^−1^, while other strains showed a biomass growth between 1.2 and 1.9 g L^−1^. Considering these growth results, the six previously mentioned strains were used for subsequent studies.

### Characterization of pesticide-tolerant bacteria

The strains selected for their tolerance and ability to grow in the presence of pesticides were characterized based on some phenotypic and biochemical characteristics (Table 2). According to Gram-staining analyses, all strains were Gram-negative, except strain C8. Most isolates exhibited cream-colored colonies; strain C8 was dark-cream colored, and strain C7 had white colonies. The morphological characteristics of the bacteria were evaluated by means of SEM. The presented micrographs in Fig. 1 show three different strains with representative morphological cell structures. Five of the six selected strains were bacillus, while strain C8 presented a coccus shape. In general, strains C4 and C10 with a bacillus cell shape and size ranging from 0.82 × 2.35 µm to 0.91 × 1.84 µm were observed, respectively. All other strains were omitted due to similarities in size. For strain C8 with a coccus cell shape, the sizes ranged from 0.76 to 1.26 µm in diameter.

**Fig. 1.**
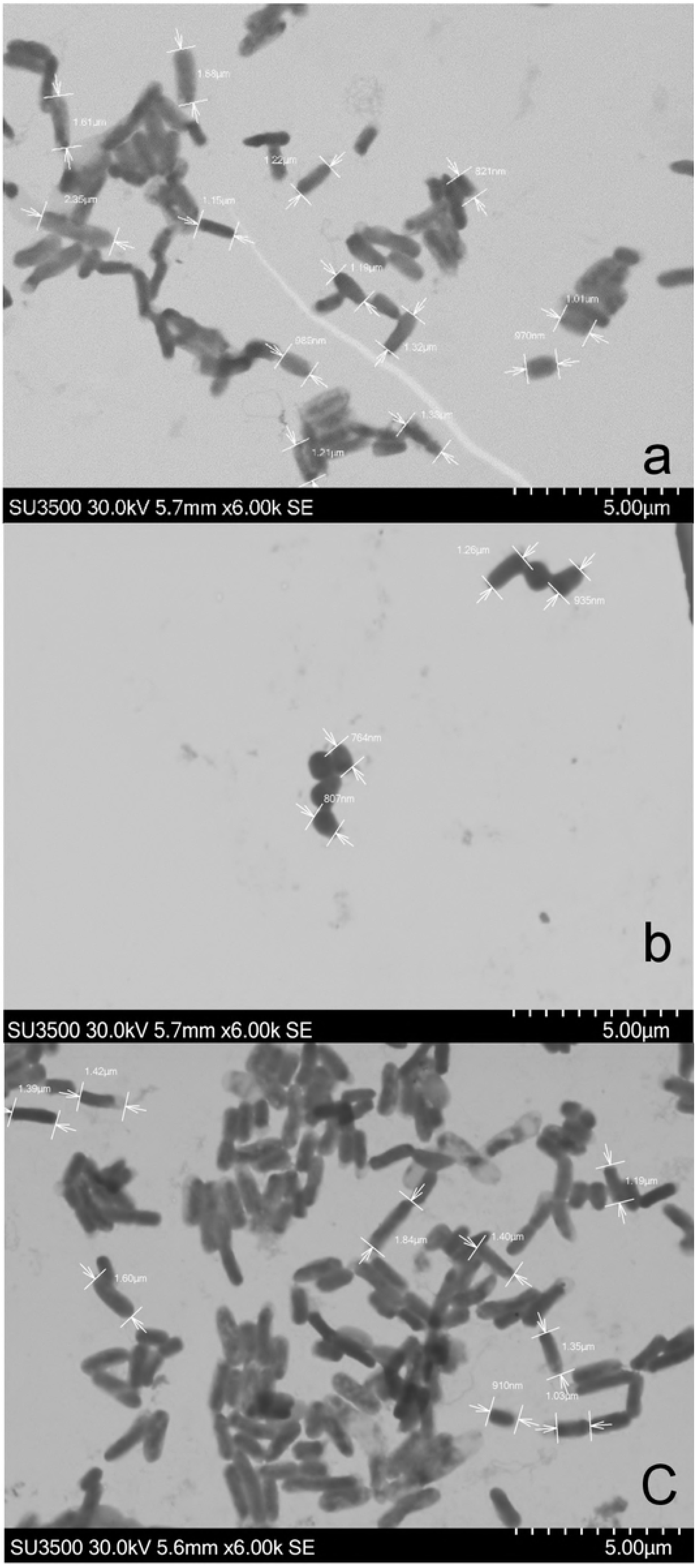
Electron scan micrographs of cells morphology of C4 (a), C8 (b) and C10 (c) strains isolated by enrichment from a biomixture of a biopurification system treated repeatedly with pesticides.

The results of biochemical and enzymatic characterization using API ZYM showed that all isolates (100%) were positive for esterase (C4), leucine aminopeptidase, acid phosphatase, and naphthol-AS-BI-phosphohydrolase. Enzymes, such as alkaline phosphatase and valine aminopeptidase, tested positive in 83% of strains, esterase lipase (C8) in 50% of strains, and lipase (C14) and trypsin in 33% strains. Strain C8 was positive for most enzymes, including α-glucosidase and β-glucosidase involved in glucose metabolism. Finally, screening of pesticide-tolerant strains isolated from the biopurification system showed that C4 and C9 strains produced five extracellular enzymes on solid R2A medium. Regarding the production of lipases and amylase, the six selected strains were positive for both enzymes, while strains C1, C4, C8, and C9 were positive for cellulolytic enzymes (Table 2).

**Table 2.** Phenotypic features and biochemical characteristics of different pesticide- tolerant bacteria isolated from biopurificaction system.

|  | (-) (+) |  | Response of strains |  |  |  |  |  |
| --- | --- | --- | --- | --- | --- | --- | --- | --- |
|  | (%) |  | C1 | C4 | C7 | C8 | C9 | C10 |
| **Color of colonies** |  |  | Cream | Cream | White | Dark-Cream | Cream | Cream |
| **Morphology** |  |  | Bacillus | Bacillus | Bacillus | Coccus | Bacillus | Bacillus |
| **Gram staining** |  |  | - | - | - | + | - | - |
| **Enzyme\*** |  |  | **C1** | **C4** | **C7** | **C8** | **C9** | **C10** |
| Control | 100 | 0 | - | - | - | - | - | - |
| Alkaline phosphatase | 16.7 | 83.3 | + | - | + | + | + | + |
| Esterase (C4) | 0 | 100 | + | + | + | + | + | + |
| Esterase lipase (C8) | 50 | 50 | - | + | - | + | + | - |
| Lipase (C14) | 66.7 | 33.3 | - | + | - | + | - | - |
| Leucine aminopeptidase | 0 | 100 | + | + | + | + | + | + |
| Valine aminopeptidase | 16.7 | 83.3 | + | + | - | + | + | + |
| Cystine aminopeptidase | 83.3 | 16.7 | - | - | - | + | - | - |
| Trypsin | 66.7 | 33.3 | - | + | - | - | + | - |
| α- Chymotrypsin | 83.3 | 16.7 | - | - | - | + | - | - |
| Acid phosphatase | 0 | 100 | + | + | + | + | + | + |
| Naphthol-AS-BI-phosphohydrolase | 0 | 100 | + | + | + | + | + | + |
| α-Galactosidase | 100 | 0 | - | - | - | - | - | - |
| β- Galactosidase | 100 | 0 | - | - | - | - | - | - |
| β-Glucuronidase | 100 | 0 | - | - | - | - | - | - |
| α- Glucosidase | 83.3 | 16.7 | - | - | - | + | - | - |
| β- Glucosidase | 83.3 | 16.7 | - | - | - | + | - | - |
| N-acetyl-β- Glucosaminidase | 100 | 0 | - | - | - | - | - | - |
| α- Mannosidase | 100 | 0 | - | - | - | - | - | - |
| α- Fucosidase | 100 | 0 | - | - | - | - | - | - |
| **Extracellular hydrolase activity #** |  |  | **C1** | **C4** | **C7** | **C8** | **C9** | **C10** |
| Amylolytic (starch 0.4%) | 0 | 100 | + | + | + | + | + | + |
| Cellulolytic (CMC 0.4%) | 33.3 | 66.7 | + | + | - | + | + | - |
| Lipolytic (tween 80%) | 0 | 100 | + | + | + | + | + | + |
| Proteolytic (milk 30%) | 66.7 | 33.3 | - | + | - | - | + | - |
| Proteolytic (gelatin 1%) | 66.7 | 33.3 | - | + | - | - | + | - |

+ : Positive reaction, - : Negative reaction; \*Analysed by API ZYM kit; # Tested by tested on R2A agar.

### Molecular and proteomic identification of bacteria

The identification of selected strains made by both 16S rDNA sequencing and MALDI-TOF/TOF MS showed similar results. The strains selected for their tolerance and ability to grow in the presence of pesticides CHL and IPR were identified based on 16S rDNA sequence analysis as bacteria belonging to the phylum *Proteobacteria*, family *Alcaligenaceae*, genus *Achromobacter* (strains C1, C7, and C10), and family *Pseudomonadaceae*, genus *Pseudomonas* (strains C4 and C9). Moreover, the phylum *Actinobacteria*, family *Nocardiaceae*, and genus *Rodococcus* (strains C8) was identified. A comparison of the 16S rDNA sequences (entire sequence compared with available sequences in GenBank) of strains C1, C4, C7, C8, C9, and C10 showed ≥ 96% similarity to those of *Achromobacter spanius*, *Pseudomonas rhodesiae*, *Achromobacter deleyi*, *Rhodococcus jialingiae*, *Pseudomonas marginalis*, and *Achromobacter kerstersii*, respectively (Table 3). To identify the phylogeny of the isolates, strains from different genera were chosen to construct the phylogenetic tree. Phylogenetic analysis (Fig. 2) based on the 16S rDNA using MEGA7 software indicated that the isolates had higher similarity with the 16S rDNA sequence from pesticide-degrading bacteria, i.e., *Pseudomonas caspiana* (strains C4 and C9), *Rhodococcus jialingiae* (strain C8), and *Achromobacter spirinitus* (C1, C7 and C10).

**Fig. 2.**
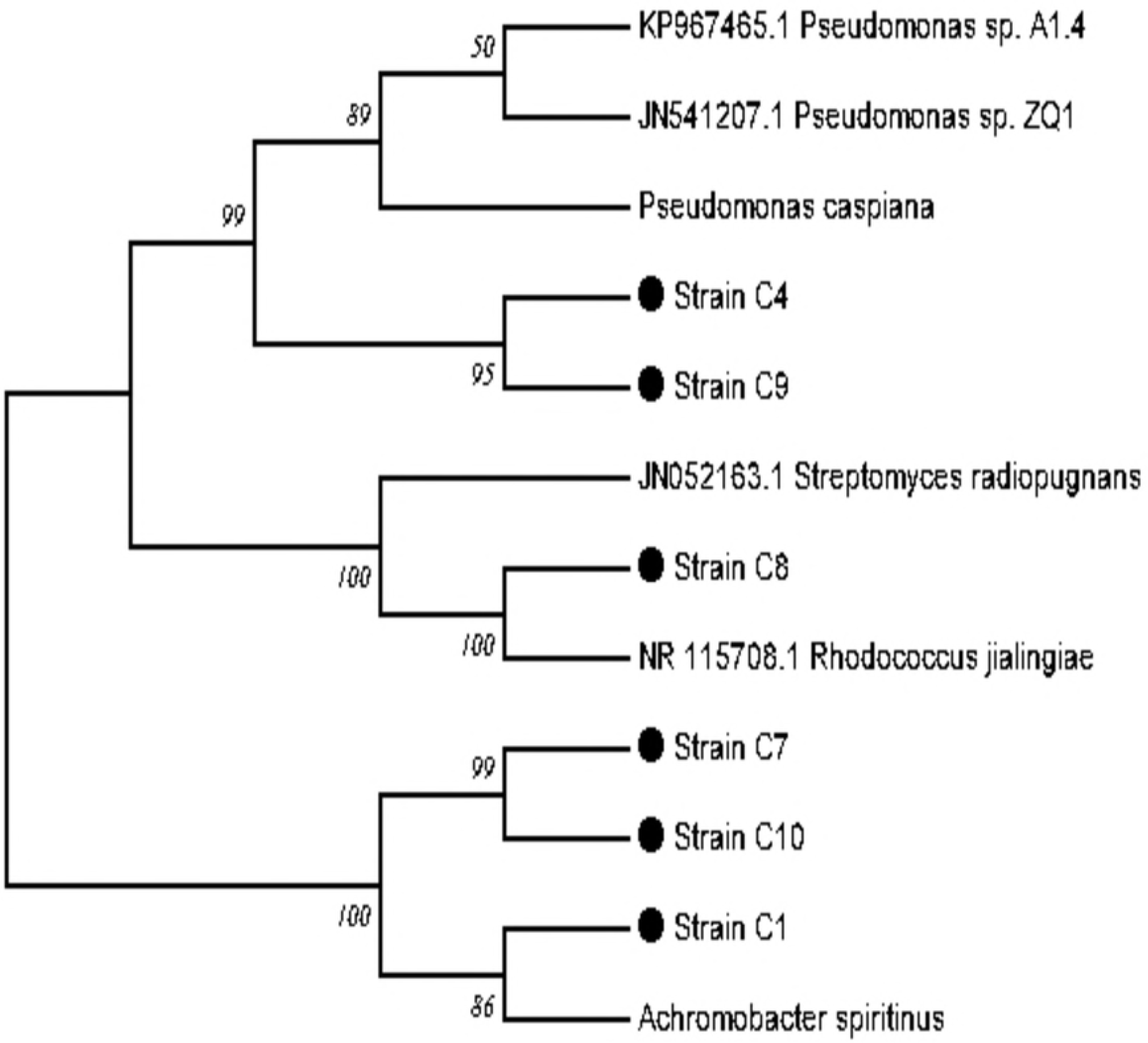
Phylogenetic tree constructed by the neighbor-joining method based on 16S rDNA sequences of studied C1, C4, C7, C8, C9, C10 strains and related ones.

**Table 3.** Phylogenetic assignment of isolated strains tolerant to chlorpyrifos (CHL) and iprodione (IPR) and their best match results with 16S rDNA gene sequences and MALDI -TOF TOF BioTyper.

| Strains | Most closely related strain<br>(NCBI accession no.)<sup>a</sup> | Identity<br>(%) | Acession n<sup>o</sup> | Identification<br>MALDI Biotyper<br>database | Score<sup>b</sup> |
| --- | --- | --- | --- | --- | --- |
| C1 | *Achromobacter spanius*<br>(MF624722.1) | 96 | MK110041 | *Achromobacter* sp. | 2.36 |
| C4 | *Pseudomonas rhodesiae*<br>(R_024911.1) | 99 | MK110043 | *Pseudomonas* sp. | 2.31 |
| C7 | *Achromobacter deleyi*<br>(NR_152014.1) | 97 | MK110044 | *Achromobacter* sp. | 2.06 |
| C8 | *Rhodococcus jialingiae*<br>(NR_115708.1) | 98 | MK110045 | *Rhodococcus* sp. | 2.54 |
| C9 | *Pseudomonas marginalis*<br>(NR_117821.1) | 99 | MK110046 | *Pseudomonas* sp. | 2.14 |
| C10 | *Achromobacter kerstersii*<br>(NR_152015.1) | 98 | MK110047 | *Achromobacter* sp. | 2.04 |

(a) Based on partial sequencing of 16S rDNA gene and comparison with those present in GenBank database from National Center for Biotechnology Information (NCBI) by using BLAST.
(b) Log score value of the MALDI-TOF MS identification.

Direct analysis of intact cells by MALDI-TOF/TOF MS showed a very good spectral quality with score identification of 2.04 to 2.54 (Table 3), safely allowing accurate identification to the genus level. Genus identification of the different strains was in agreement with the 16S rDNA sequence identification. The dendrogram constructed using the MALDI Biotyper data of the six bacteria in the presence of CHL and IPR showed that *Achromobacter* sp. strains C1, C7, and C10 were differentiated and grouped separately when exposed to different pesticides, and a similar response was observed for *Pseudomonas* sp. strains C4 and C9 (Fig. 3).

**Fig. 3.**
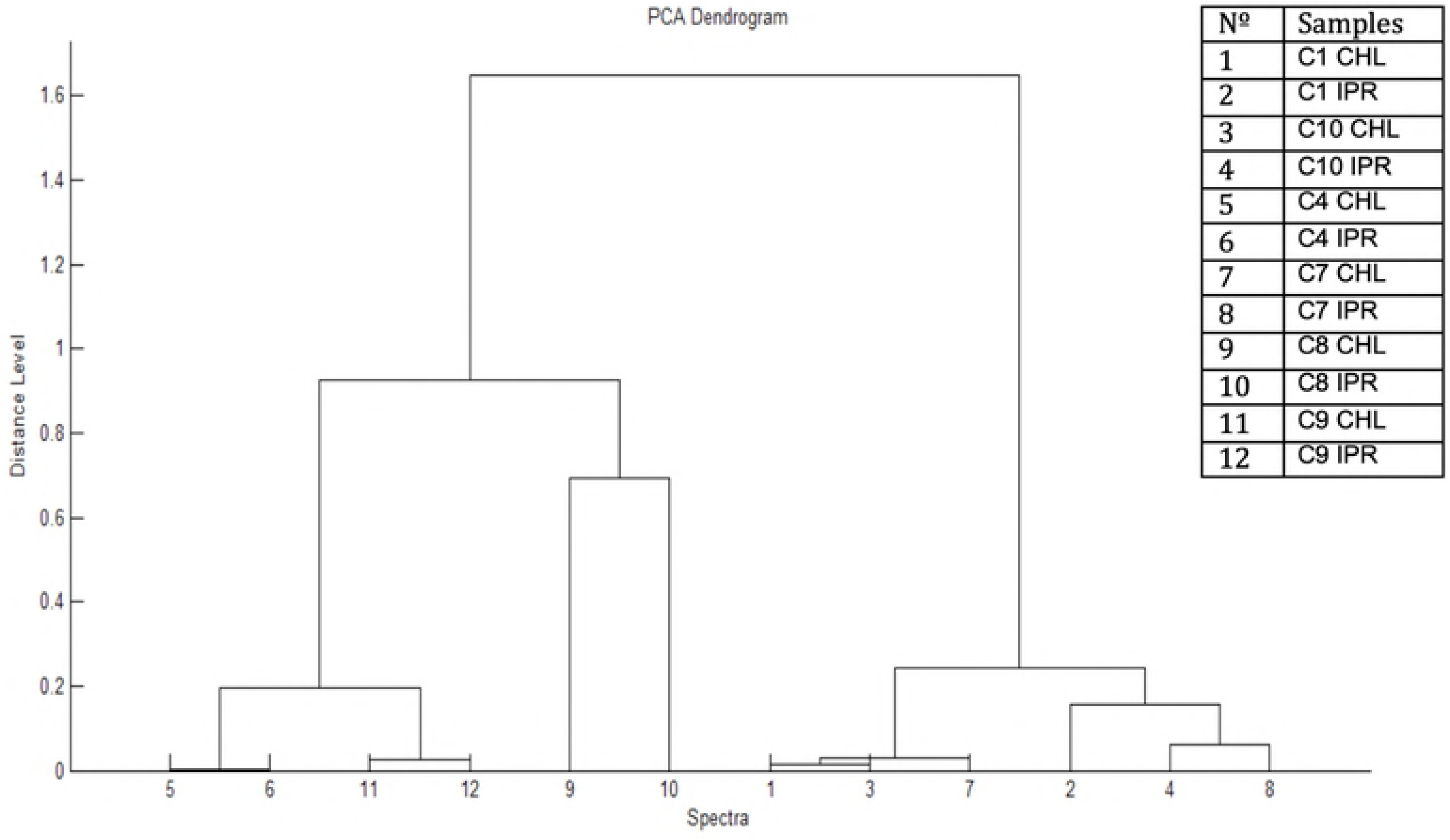
Dendrogram obtained by MALDI Biotyper Compass 4.1 software (Bruker Daltonics, Bremen, Germany) of C1, C4, C7, C8, C9, C10 strains after enrichment with CHL and IPR of liquid cultures.

### Growth and degradation of pesticides in liquid cultures

Biomass growth of the six tolerant-pesticides strains was evaluated at different incubation times and increasing pesticide concentrations, observing that bacterial growth decreased proportionally (R^2^ > 0.96) as both pesticide concentrations increased from 10 to 100 mg L^−1^. As observed in Fig. 4, all bacteria exposed to CHL concentrations from 10 to 50 mg L^−1^ showed an increase in biomass over time. However, a high inhibition of biomass growth was observed in all strains cultivated in liquid medium supplemented with 100 mg L^−1^ CHL. In the same way, only *Achromobacter* sp. strain C1 and *Pseudomonas* sp. strain C4 showed the highest tolerance to 100 mg L^−1^ of CHL, which was compared to microbial growth observed in the control treatment without pesticide. In general, the µ max for the studied strains ranged from 0.18 to 0.48 h^−1^ in the treatment without pesticides, and these values decreased showing a µ rate between 0.02 to 0.16 h^−1^ for CHL added at the highest concentration. *Pseudomonas* sp. strain C4 showed to be the most tolerant strain to CHL in relation to the growth observed in the control treatment (Table 4).

**Fig. 4.**
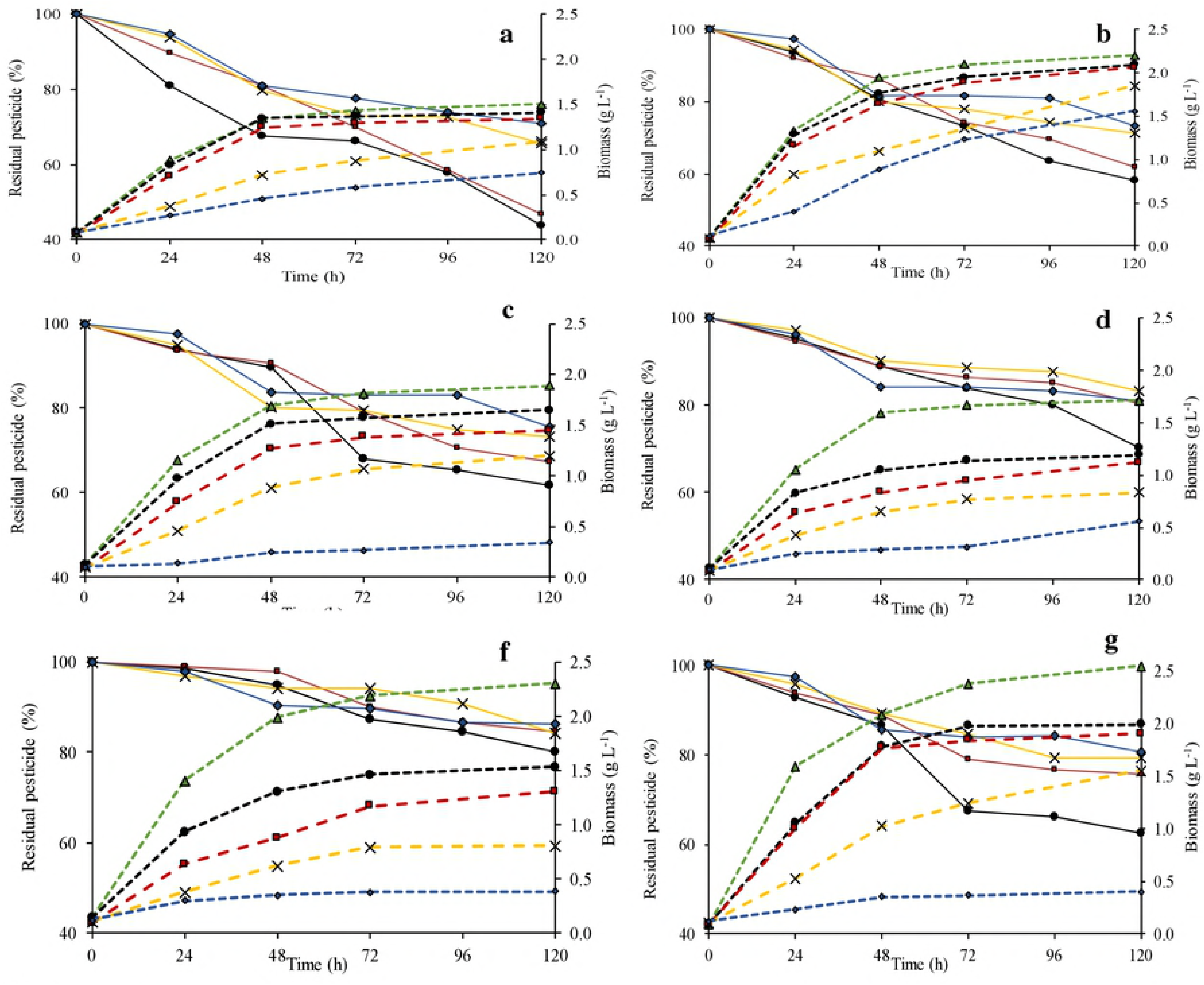
Remaining chlorpyrifos (CHL) (%) and biomass by C1 (a), C4 (b), C7 (c), C8 (d), C9 (3), C10 (f) strains at initial CHL concentration of 0, 10, 20, 50 and 100 mg L^−1^, evaluated during 120 h. 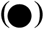 CHL 10 mg L^−1^; 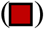 CHL 20 mg L^−1^; (X) CHL 50 mg L^−1^; 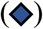 CHL 100 mg L^−1^; 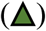 Control without CHL. Continuous line: Remaining CHL (%); Dotted line: biomass (g L^−1^).

**Table 4.** First-order kinetics parameter for chlorpyrifos (CHL) and iprodione (IPR) removal and specific growth rate (µ) of strains C1, C4, C7, C8, C9 and C,10 in liquid medium supplemented with 0-100 mg L^−1^ of pesticide individually.

| Chlorpyrifos |  |  |  |  |  |  |  |  |  |  |  |  |  |  |  |  |
| --- | --- | --- | --- | --- | --- | --- | --- | --- | --- | --- | --- | --- | --- | --- | --- | --- |
| Strain | 10 mg L<sup>-1</sup> |  |  |  | 20 mg L<sup>-1</sup> |  |  |  | 50 mg L<sup>-1</sup> |  |  |  | 100 mg L<sup>-1</sup> |  |  |  |
| | $\mu$<br>max<br>(h<sup>-1</sup>) | R<br>(%) | *k*<br>(d<sup>-1</sup>) | T<sub>1/2</sub><br>(d) | $\mu$<br>max<br>(h<sup>-1</sup>) | R<br>(%) | *k*<br>(d<sup>-1</sup>) | T<sub>1/2</sub><br>(d) | $\mu$<br>max<br>(h<sup>-1</sup>) | R<br>(%) | *k*<br>(d<sup>-1</sup>) | T<sub>1/2</sub><br>(d) | $\mu$<br>max<br>(h<sup>-1</sup>) | R<br>(%) | *k*<br>(d<sup>-1</sup>) | T<sub>1/2</sub><br>(d) |
| C1 | 0.17 | 56.25±0.10 a | 0.147 | 4.7 | 0.16 | 53.11±0.01 a | 0.149 | 4.6 | 0.11 | 34.46±0.02 a | 0.085 | 8.2 | 0.09 | 29.10±0.01 a | 0.072 | 9.7 |
| C4 | 0.30 | 41.80±0.02 b | 0.113 | 6.1 | 0.29 | 38.04±0.04 b | 0.097 | 7.2 | 0.25 | 28.60±0.09 b | 0.069 | 10.0 | 0.16 | 28.04±0.01 a | 0.061 | 11.4 |
| C7 | 0.26 | 38.34±0.02 c | 0.108 | 6.4 | 0.22 | 32.56±0.00 b | 0.084 | 8.2 | 0.16 | 26.89±0.07 c | 0.066 | 10.6 | 0.02 | 24.34±0.07 b | 0.054 | 12.9 |
| C8 | 0.22 | 29.75±0.04 d | 0.067 | 10.4 | 0.19 | 19.70±0.01 cd | 0.041 | 16.8 | 0.15 | 16.89±0.02 e | 0.036 | 19.4 | 0.08 | 18.98±0.04 c | 0.042 | 16.4 |
| C9 | 0.28 | 19.82±0.04 e | 0.047 | 14.7 | 0.22 | 15.29±0.05 d | 0.037 | 18.6 | 0.14 | 15.88±0.02 f | 0.030 | 22.8 | 0.11 | 13.57±0.05 d | 0.032 | 21.9 |
| C10 | 0.40 | 37.45±0.01 c | 0.105 | 6.6 | 0.39 | 24.26±0.02 c | 0.060 | 11.5 | 0.27 | 20.52±0.04 d | 0.050 | 13.8 | 0.14 | 19.46±0.02 c | 0.044 | 15.8 |

| Iprodione |  |  |  |  |  |  |  |  |  |  |  |  |  |  |  |  |
| --- | --- | --- | --- | --- | --- | --- | --- | --- | --- | --- | --- | --- | --- | --- | --- | --- |
| Strain | 10 mg L<sup>-1</sup> |  |  |  | 20 mg L<sup>-1</sup> |  |  |  | 50 mg L<sup>-1</sup> |  |  |  | 100 mg L<sup>-1</sup> |  |  |  |
| | $\mu$<br>max<br>(h<sup>-1</sup>) | R<br>(%) | *k*<br>(h<sup>-1</sup>) | T<sub>1/2</sub><br>(h) | $\mu$<br>max<br>(h<sup>-1</sup>) | R<br>(%) | *k*<br>(h<sup>-1</sup>) | T<sub>1/2</sub><br>(h) | $\mu$<br>max<br>(h<sup>-1</sup>) | R<br>(%) | *k*<br>(h) | T<sub>1/2</sub><br>(h) | $\mu$<br>max<br>(h<sup>-1</sup>) | R<br>(%) | *k*<br>(h<sup>-1</sup>) | T<sub>1/2</sub><br>(h) |
| C1 | 0.18 | 97.60±0.07 b | 0.160 | 4 | 0.17 | 94.40±0.01 b | 0.123 | 6 | 0.15 | 90.30±0.15 b | 0.099 | 7 | 0.11 | 84.30±0.00 b | 0.078 | 9 |
| C4 | 0.26 | 94.20±0.02 c | 0.121 | 6 | 0.23 | 92.70±0.00 c | 0.112 | 6 | 0.22 | 87.50±0.00 c | 0.089 | 8 | 0.09 | 81.50±0.14 c | 0.072 | 10 |
| C7 | 0.25 | 87.10±0.09 e | 0.087 | 8 | 0.20 | 84.70±0.31 d | 0.079 | 9 | 0.17 | 79.60±0.17 e | 0.068 | 10 | 0.05 | 76.60±0.25 e | 0.062 | 11 |
| C8 | 0.24 | 88.60±0.00 d | 0.092 | 8 | 0.22 | 85.60±0.15 d | 0.082 | 8 | 0.20 | 81.10±0.05 d | 0.071 | 10 | 0.12 | 77.80±0.10 d | 0.065 | 11 |
| C9 | 0.35 | 98.90±0.00 a | 0.193 | 4 | 0.34 | 96.90±0.00 a | 0.148 | 5 | 0.32 | 93.10±0.09 a | 0.113 | 6 | 0.22 | 91.20±0.02 a | 0.103 | 7 |
| C10 | 0.43 | 81.10±0.70 f | 0.070 | 10 | 0.42 | 77.10±0.01 e | 0.062 | 11 | 0.36 | 74.20±0.09 f | 0.058 | 12 | 0.24 | 70.40±0.18 f | 0.052 | 13 |

Values of removal (% R) within a concentration with the same letter are not significantly different based on the Tukey test ( $\alpha=0.05$ ), the average values and the standard error are presented (n= 3); R (%): removal of pesticides, *k*: rate constant, T<sub>1/2</sub>: half-life time, $\mu$ : specific growth rate.

Similar to that observed for CHL, biomass of bacteria exposed to IPR increased over time, up to 50 mg L^−1^ IPR concentration, where growth decreased as the pesticide concentration increased (Fig. 5). In the control treatments, a biomass between 1.43 and 2.38 g L^−1^ and µ max from 0.18 to 0.48 h^−1^ were observed, instead of biomass between 0.78 and 1.80 g L^−1^ and µ max from 0.15 to 0.36 h^−1^ at 50 mg L^−1^ of IPR. Application of 100 mg L^−1^ IPR in the liquid medium caused a marked inhibition of microbial growth with biomass ranging between 0.34 to 1.06 g L^−1^ and a µ max between 0.05 and 0.24 h^−1^. *Achromobacter* sp. strain C1 and *Pseudomonas* sp. strain C9 were the most tolerant strains to IPR in relation to the growth observed in the control treatment (Table 4).

**Fig. 5.**
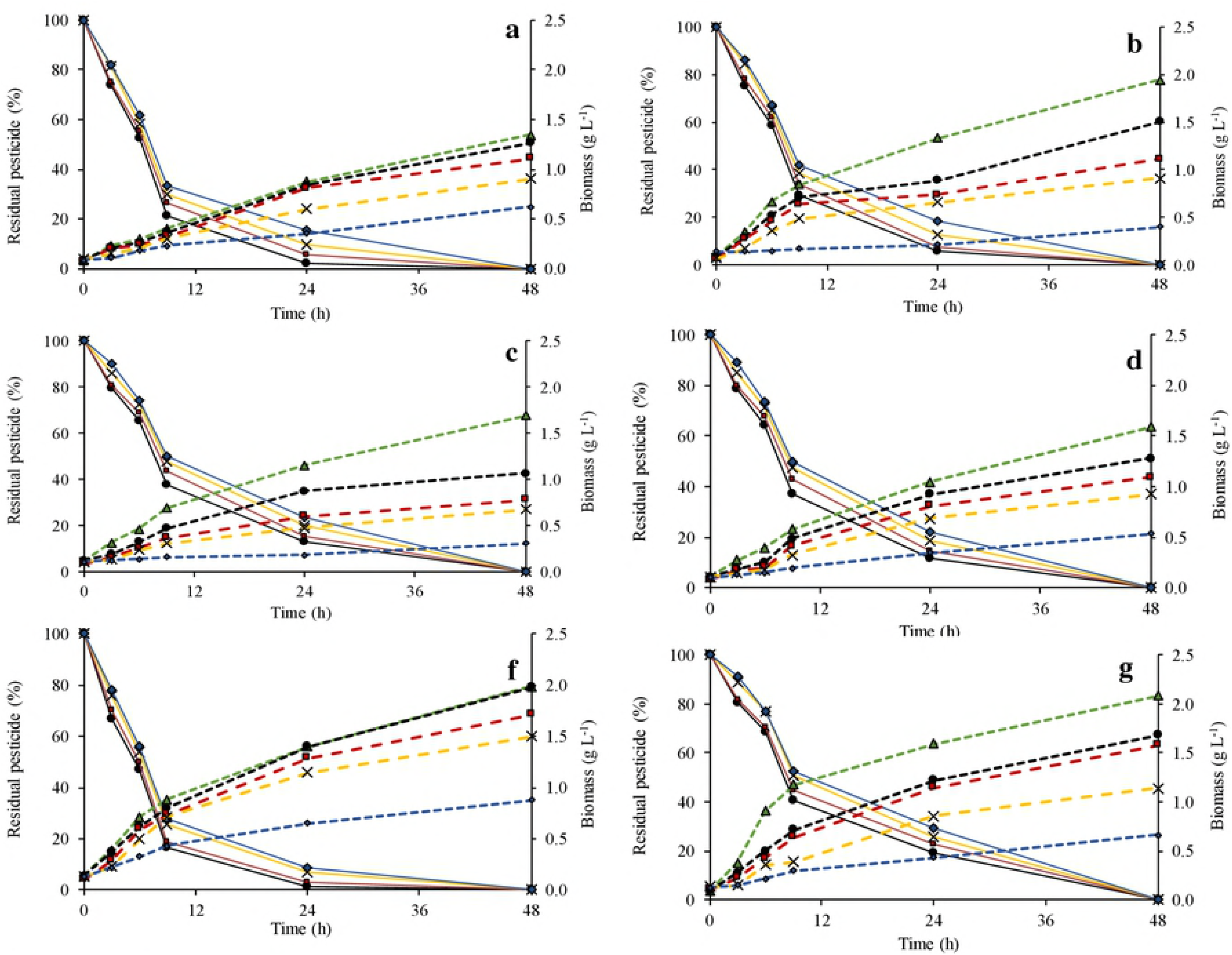
Remaining iprodione (IPR) (%) and biomass by C1 (a), C4 (b), C7 (c), C8 (d), C9 (3), C10 (f) strains at initial CHL concentration of 0, 10, 20, 50 and 100 mg L^−1^, evaluated during 120 h. 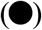 CHL10 mg L^−1^; 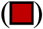 CHL 20 mg L^−1^; (x) CHL 50 mg L^−1^; 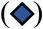 CHL 100 mg L^−1^; 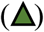 Control without CHL. Continuous line: Remaining IPR (%); Dotted line: biomass (g L^−1^).

With respect to CHL removal, it decreased for all bacteria strains as the contaminant concentration increased from 10 to 100 mg L^−1^. *Achromobacter* sp. strain C1 showed the best CHL removal (56–29%) after 120 h of incubation, which was significant (p ≤ 0.05) relative to that by the other five strains for all CHL concentrations. In this context, the kinetic data showed that CHL removal by the C1 strains were characterized by a rate constant of 0.147–0.072 d^−1^ and T_1/2_ 4.7–9.7 d^−1^ in liquid medium treated with 10 and 100 mg L^−1^ CHL. This trend was closely followed by the *Pseudomonas* sp. strains C4, *Achromobacter* sp. strains C7, and *Achromobacter* sp. strain C10 with a removal between 42–27%, 38–24%, and 37–19%, respectively. However, the lowest degradation was observed for the *Pseudomonas* sp. strain C9 with CHL degradation between 19–13%, a rate constant between 0.030–0.047 d^−1^, and T_1/2_ 14–22 d^−1^ (Table 4).

With respect to IPR degradation, when concentrations increased from 10 to 100 mg L^−1^, it was removed efficiently (81–98%) by all strains, requiring only 24 h of incubation. IPR was not detected at 48 h. The significantly (p ≤ 0.05) highest IPR removal (between 91.2–98.9%) was observed for the *Pseudomonas* sp. strain C9 relative to that for the other strains at all IPR concentrations. According to the kinetic parameters, strain C9 showed the highest rate constant of 0.193 h^−1^ for IPR added at 10 mg L^−1^ and a maximum T_1/2_ of 7 h^−1^ when IPR was added at a concentration of 100 mg L^−1^. For *Achromobacter* sp. strain C10, the lowest IPR removal (70–81%) was observed after 24 h of incubation and a T_1/2_ ranging from 11 to 13 h^−1^ (Table 4).

In parallel to pesticides removal from the liquid medium, the concentrations of metabolite TCP and 3,5-DCA produced during CHL and IPR removal, respectively, were analyzed (Table 5).

**Table 5.** Production of 3,5,6-trichlo-2-pyridinol (TCP) and 3,5-DCA by strains C1, C4, C7, C8, C9 and C,10 in liquid medium supplemented with 0-100 mg L^−1^ of chlorpyrifos (CHL) and iprodione (IPR) individually.

| Strain<br>s | Time<br>(h) | Chlorpyrifos |  |  |  | Iprodione |  |  |  |
| --- | --- | --- | --- | --- | --- | --- | --- | --- | --- |
|  |  | 10 mg L<sup>-1</sup> | 20 mg L<sup>-1</sup> | 50 mg L<sup>-1</sup> | 100 mg L<sup>-1</sup> | 10 mg L<sup>-1</sup> | 20 mg L<sup>-1</sup> | 50 mg L<sup>-1</sup> | 100 mg L<sup>-1</sup> |
|  |  | Metabolite | Metabolite | Metabolite | Metabolite | Metabolite | Metabolite | Metabolite | Metabolite |
|  |  | (mg L<sup>-1</sup>) | (mg L<sup>-1</sup>) | (mg L<sup>-1</sup>) | (mg L<sup>-1</sup>) | (mg L<sup>-1</sup>) | (mg L<sup>-1</sup>) | (mg L<sup>-1</sup>) | (mg L<sup>-1</sup>) |
| C1 | 24 | 0.17±0.01 | 0.36±0.02 | 0.82±0.02 | 1.61±0.02 | 0.21±0.00 | 0.35±0.01 | 0.29±0.01 | 0.47±0.01 |
|  | 72 | 0.22±0.29 | 0.41±0.01 | 0.94±0.01 | 1.84±0.03 | 0.24±0.05 | 0.36±0.02 | 0.45±0.02 | 0.65±0.01 |
|  | 120 | 0.50±0.02 | 0.69±0.01 | 1.62±0.02 | 2.10±0.02 | 0.38±0.00 | 0.42±0.04 | 0.52±0.06 | 0.87±0.10 |
| C4 | 24 | 0.12±0.02 | 0.32±0.02 | 0.80±0.02 | 1.40±0.02 | 0.15±0.03 | 0.29±0.01 | 0.26±0.01 | 0.35±0.01 |
|  | 72 | 0.14±0.01 | 0.33±0.01 | 0.88±0.04 | 1.47±0.04 | 0.20±0.06 | 0.34±0.01 | 0.41±0.06 | 0.59±0.06 |
|  | 120 | 0.34±0.01 | 0.52±0.00 | 1.48±0.08 | 1.50±0.04 | 0.34±0.01 | 0.41±0.07 | 0.49±0.02 | 0.86±0.06 |
| C7 | 24 | 0.08±0.02 | 0.28±0.02 | 0.74±0.02 | 1.42±0.02 | 0.32±0.03 | 0.16±0.03 | 0.19±0.07 | 0.27±0.05 |
|  | 72 | 0.10±0.00 | 0.30±0.01 | 0.77±0.02 | 1.46±0.04 | 0.18±0.02 | 0.26±0.04 | 0.29±0.04 | 0.41±0.07 |
|  | 120 | 0.29±0.01 | 0.48±0.02 | 1.27±0.04 | 1.47±0.02 | 0.30±0.02 | 0.35±0.01 | 0.37±0.02 | 0.75±0.13 |
| C8 | 24 | 0.08±0.02 | 0.27±0.02 | 0.66±0.02 | 1.38±0.02 | 0.15±0.02 | 0.20±0.03 | 0.22±0.04 | 0.35±0.01 |
|  | 72 | 0.07±0.01 | 0.26±0.01 | 0.73±0.04 | 1.46±0.03 | 0.20±0.00 | 0.32±0.05 | 0.35±0.05 | 0.50±0.02 |
|  | 120 | 0.24±0.00 | 0.42±0.02 | 0.96±0.03 | 1.45±0.00 | 0.31±0.04 | 0.39±0.01 | 0.42±0.04 | 0.85±0.08 |
| C9 | 24 | 0.07±0.02 | 0.26±0.02 | 0.61±0.02 | 1.36±0.03 | 0.25±0.00 | 0.38±0.03 | 0.37±0.03 | 0.47±0.08 |
|  | 72 | 0.06±0.01 | 0.25±0.01 | 0.78±0.04 | 1.38±0.04 | 0.26±0.02 | 0.38±0.01 | 0.47±0.04 | 0.70±0.04 |
|  | 120 | 0.21±0.00 | 0.47±0.01 | 0.91±0.01 | 1.40±0.00 | 0.31±0.00 | 0.49±0.01 | 0.59±0.03 | 0.95±0.00 |
| C10 | 24 | 0.06±0.02 | 0.25±0.02 | 0.74±0.02 | 1.56±0.02 | 0.07±0.01 | 0.10±0.02 | 0.10±0.00 | 0.22±0.01 |
|  | 96 | 0.07±0.03 | 0.26±0.02 | 0.83±0.01 | 1.45±0.08 | 0.17±0.06 | 0.12±0.04 | 0.22±0.03 | 0.39±0.02 |
|  | 120 | 0.28±0.00 | 0.47±0.01 | 0.98±0.01 | 1.38±0.00 | 0.21±0.05 | 0.25±0.00 | 0.35±0.00 | 0.65±0.05 |

The average values and the standard error are presented (n= 3)

The results showed that over time and when the pesticide concentration increased, the metabolite concentrations increased. TCP production was highest when liquid media was treated with *Achromobacter* sp. strain C1, detecting concentrations between 0.504–2.098 mg L^−1^ after 120 h of incubation. For all other strains, TCP concentrations ranged from 0.214 mg L^−1^ produced by *Pseudomonas* sp. strain C9 during the treatment of 10 mg L^−1^ CHL to 1.498 mg L^−1^ produced by *Pseudomonas* sp. strains C4 during the treatment of 100 mg L^−1^ CHL. In relation to IPR degradation, the results showed that the metabolite began to appear in the liquid medium at 9 h of incubation. After 48 h of incubation, 3,5-DCA concentrations ranged between 0.210–0.384 mg L^−1^ and 0.648–0.945 mg L^−1^ after treatment of 10 and 100 mg L^−1^ IPR, respectively. Moreover, the highest 3,5-DCA concentrations were produced during the treatment of 20, 50, and 100 mg L^−1^ IPR by *Pseudomonas* sp. strain C9.

## Discussion

Inappropriate pesticide management has resulted in their release into the environment, including food. Therefore, efforts to develop technologies that guarantee effective treatment of pesticide residues have been made. Due to the use of microorganisms in pesticide treatment, it is extremely important to previously determine their potential for removal from liquid media under optimal conditions. Screening of degrading microorganisms through an enrichment procedure in the pesticides-contaminated system allows for selection of potential isolates with a high tolerance and maximal degrading activity [22]. In our study, the microorganism isolation matrix consisted of the biopurification system repeatedly treating a biomixture of pesticides. Pesticide application in the biomixture did not cause relevant effects on total cultivable bacteria; however, bacteria are active, which is associated with the high efficiency of pesticides degradation shown in this biopurification system [15,32]. In this context, the use of a biomixture allowed us to obtain ten tolerant strains. However, six strains were selected for their high tolerance and ability to grow in the presence of CHL and IPR; both contaminants were demonstrated as carbon and energy sources presumably via partial transformation reactions that can occur with different chemical classes of pesticides.

The selected strains were characterized in different ways. Enzymatic characterization using the ApiZym test showed production of different enzymes for each substrate. It should be noted that some authors have highlighted bacteria-produced enzymes obtained from contaminated sites of high biotechnological, clinical, and industrial interest [26,29]. In our study, strains C4, C8, and C9 presented the highest biochemical activity, being positive for at least 13 of the 24 enzymes tested.

Tolerant and characterized strains were identified using two methods: 16S rDNA gene sequencing, which is widely used to determine microorganism taxonomic positions, and MALDI-TOF/TOF MS, which identifies and classifies an organism according to the spectral profile of its ribosomal proteins. According to our results, both identifications were concordant. The phylogenetic analysis of isolates showed a closer relation with bacteria from genera *Pseudomonas*, *Achromobacter*, and *Rhodococcus*, known as metabolically active microorganisms capable of degrading many pesticides, including CHL and IPR [24,27]. Use of MALDI-TOF/TOF MS has led to a new era of routine and rapid identification of different organisms, including environmental bacteria [11]. In our study, dendrograms generated by MALDI Biotyper were able to separate or assemble different strains in the same cluster, depending on pesticides exposure, which could explain effects on proteins due to environmental conditions [33].

The results of bacterial growth cultured in the presence of CHL and IPR in liquid medium demonstrated that pesticides produced an inhibitory effect at concentrations above 50 mg L^−1^. Although various researchers have reported that CHL and IPR in liquid media could be used as a source of C and energy for growth [24,25], metabolites, which are more toxic than the parent compound and with antimicrobial properties to inhibit microbial growth, can be produced during microbial treatment [34]. IPR is known to inhibit DNA and RNA synthesis, cell division, and cellular metabolism in fungi; however, there is limited information that IPR may inhibit environmental microbes [35]. According to our results, *Achromobacter* sp. C1 and *Pseudomonas* sp. C4 were the most tolerant strains to CHL, and *Achromobacter* sp. C1 and *Pseudomonas* sp. C9 were tolerant to IPR, which was associated with a minor difference in growth relative to that for the control treatment. Consistently, both strains presented the highest removal of pesticides. *Achromobacter* and *Pseudomonas* genera are both microbial groups recognized by their ability to remove pesticides, e.g. *Achromobacter xylosoxidans* strain CS5 removes endosulfan [36], *Arthrobacter* sp. BS2 and *Achromobacter* sp. SP1 degrade diuron and their metabolite 3,4-dichloroaniline [37], *Pseudomonas* sp. and *Achromobacter* sp. isolated from agricultural soil degrade atrazine [38], *Arthrobacter* sp. strain C1 and *Achromobacter* sp. strain C2 isolated from soil degrade IPR [25], and *Pseudomonas* spp. has been described as a CHL-degrader [24]. The highest CHL removal by *Achromobacter* sp. strain C1 could be explained by the presence or activity of the enzyme alkaline phosphatase, as this enzyme is a phosphomonoesterase that regulates CHL degradation through hydrolysis of O-P bonds [39]. Similarly, the presence of diverse enzymes in *Pseudomonas* sp. strain C9 could influence fast degradation, and therefore reduce T_1/2_ required for pesticide reduction.

Previous researchers have reported that CHL removal by bacteria occurred through formation of metabolites, such as CHL-oxon, 3,5,6-trichloro-2-methoxypyridine, 2-chloro-6-hydroxypyridine, and TCP. We evaluated TCP as a primary metabolite at different times. The results showed that product levels slightly increased over time, reaching a maximum concentration of 2.098 mg L^−1^ in the liquid medium. According to our results, TCP was not metabolized by any strain, resulting in its accumulation in the liquid medium. Therefore, TCP accumulation and the presence of chlorine atoms on the pyridinol ring caused a toxic effect on the microorganisms [39], resulting in incomplete CHL removal in the time evaluated. Nonetheless, the removal of 10 mg L^−1^ and 20 mg L^−1^ of CHL was effectively performed by the *Achromobacter* sp. strain C1, probably requiring only few days more to achieve complete CHL elimination. A study reported that *A. xylosoxidans* JCp4 was able to mineralize 100 mg L^−1^ CHL completely after ten days with only a transient accumulation of TCP [40]. Our work constitutes one of the few reports of *Achromobacter* as CHL-degraders.

The appearance of 3,5-DCA, recognized as the major metabolite of IPR degradation, their at 9 h of incubation was observed at concentrations lower than 0.5 mg L^−1^. The appearance of 3,5-DCA was coincident with the fastest decrease of IPR levels. After this time, 3,5-DCA concentrations were slightly increased, such that no IPR residues were found after 48 h of incubation. Although IPR is a common fungicide frequently used in crops and with a classification of “probable carcinogen to humans,” treatment to eliminate IPR using microorganisms has been poorly studied. Some studies reported IPR and 3,5-DCA degradation by microorganisms isolated from soil, *Arthrobacter* sp. strains C1, and *Achromobacter* sp. strains C2 from liquid medium, showing a T_1/2_ of 2.3 h and 19.5 h, respectively [9]. In our study, a small amount of time was required for *Achromobacter* sp. strains C1 to remove 50% of the contaminant from liquid medium (T_1/2_ between 4–9 h), which might signify the environmental adaptation of this bacteria being exposed to continued pesticide application in the biomixture used for their isolation. According to Campos *et al*. [9], IPR removal could occur via initial hydrolysis to isopropylamine and metabolite I (3,5-dichlorophenyl-carboxamide) and then to metabolite II (3,5-dichlorophenylurea-acetate) before being hydrolyzed to 3,5-DCA and probably glycine. Similar results were reported by Cao *et al*. [35] for a *Microbacterium* sp. strain CQH-1 isolated via the enrichment culture technique from a soil with previous exposure to IPR.

In this study, we described different bacteria isolated from a biomixture used in a biopurification system that received continuous pesticide applications. These bacteria were capable of degrading compounds such as CHL and IPR. Given their identification and ability to remove contaminants, *Achromobacter* sp. strain C1 and *Pseudomonas* sp. strains C9 appear as promising microorganisms for treatment of matrices contaminated with CHL, IPR, or their mixture. The results of this study will help to improve current technologies for biodegradation of this commonly used insecticide and fungicide in response the problem of pesticide contamination.

## Acknowledgment

This work was supported by FONDECYT No. 1161481, GAP 2017-DIUFRO and, CONICYT/FONDAP/15130015 projects.

